# Community Detection in a Weighted Directed Hypergraph Representation of Cell-to-cell Communication Networks

**DOI:** 10.1101/2020.11.16.381566

**Authors:** Rui Hou, Michael Small, Alistair R. R. Forrest

## Abstract

Cell-to-cell communication is mainly triggered by ligand-receptor activities. Through ligand-receptor pairs, cells coordinate complex processes such as development, homeostasis, and immune response. In this work, we model the ligand-receptor-mediated cell-to-cell communication network as a weighted directed hypergraph. In this mathematical model, collaborating cell types are considered as a node community while the ligand-receptor pairs connecting them are considered a hyperedge community. We first define the community structures in a weighted directed hypergraph and develop an exact community detection method to identify these communities. We then modify approximate community detection algorithms designed for simple graphs to identify the nodes and hyperedges within each community. Application to synthetic hypergraphs with known community structure confirmed that one of the proposed approximate community identification strategies, named HyperCommunity algorithm, can effectively and precisely detect embedded communities. We then applied this strategy to two organism-wide datasets and identified putative community structures. Notably the method identifies non-overlapping edge-communities mediated by different sets of ligand-receptor pairs, however node-communities can overlap.

## 1. Introduction

A central feature of complex multicellular organisms is the establishment of multiple differentiated cell types with specialized functions during development [1, 2]. These cell types are organized into higher-level units such as niches, organs, organ systems and organisms and rely on coordinated activity of different cell-types for normal operation. Coordination of different cell types depends on chemical signals known as ligands (this term encompasses peptides, lipids, ions and other molecules), which are secreted from, or located on the surface of, broadcasting cells. Cognate receptors on target cells recognise these ligands and the binding of the ligand results in the target cell altering its state [3]. Mechanisms for cell-to-cell communication are key to multicellular coordination during embryonic development and in various metabolic and physiological processes [4–10]. Dysfunctions in signalling pathways, therefore, have been associated with a variety of diseases such as diabetes, cancer, developmental disorders and many others [11–15].

Previously Ramilowski *et al.* [16] constructed the first draft map of ligand-receptor-mediated cell-to-cell interactions across a panel of 144 human primary cell types. To identify key cell-to-cell communication paths, most contemporary studies [17–20] only consider simple graph models in which a cell-type pair can be connected by a ligand and its cognate receptor. Gene ontology enrichment analyses of communication networks modelled using simple graphs has been shown previously to identify enriched biological processes involving cells from a common lineage (e.g. receptors and ligands involved in signalling from haematological cells to haematological cells known to be involved in the immune response [21]) and between those from different lineages (e.g. signals from endothelial cells to haematological cells known to be involved in wound healing [22]).

This simple graph-based approach however drastically masks the complexity of a multicellular interaction network. The reality is that multiple cell types can express the same receptor or ligand. Thus for a defined ligand-receptor pair, multiple cell types may produce the ligand and the cognate receptor could be expressed in more than one cell type. Such a many-to-many structure in graph-theoretic terms is a directed hyperedge (from the cell type(s) expressing the ligand to the target cell type(s) expressing the receptor). Additionally, as the expression levels of the same ligand/receptor vary across different cell types, the cell-to-cell communication network involves weighted directed hyperedges that represent these distinct expression levels. To date only Tsuyuzaki *et al.* [23] have attempted to use a hypergraph model to identify key ligand-receptor-mediated cell-to-cell communications. Additionally, we have previously estimated that there are in the order of 100 ligand-receptor mediated signalling pairs between any two cell types (based on our analysis of the Functional ANnoTation Of the Mammalian genome (FANTOM5) data [1]). Hence, the network involves many weighted directed hyperedges via diverse ligand-receptor pairs. These cannot be reduced to a few representative hyperedges. The graph-based methods proposed in this paper are able to reflect these complexities.

In this paper, we first propose a weighted directed hypergraph-based mathematical description of the ligand-receptor mediated cell-to-cell interaction network. For a ligand-receptor pair, our hyperedge model explicitly characterizes the weighted many-to-many relationship. From this model, we next define the community structure in the directed hypergraph, with the assumption that cell types that participate in the same biological process would exhibit a high degree of clustering. In order to identify the defined communities, we design both exact and approximate methods. Since the community structures in the real biological samples are unknown, we tested two proposed approximate detection methods on synthetic weighted directed hypergraphs with pre-defined communities generated by our benchmark generation framework. The results show that our approximate algorithms robustly detect these pre-defined communities and that the methods scale well as the input data and number of embedded communities is increased. Lastly, when applied to organism-wide datasets, the methods identify biologically plausible communities of cell-types suggesting our methods have utility in identifying strongly communicating cell types in real cell-to-cell communication networks.

## 2. Methods

### 2.1 Hypergraph construction and biological background

The definitions below, concerning weighted directed hypergraphs, are from [24, 25]. The concept of this kind of hypergraph generalizes the usual directed hypergraph [26, 27].

#### Definition 1.

(Weighted directed hypergraph) A weighted directed hypergraph is a tuple *C*(*V, E, W, f*) with *V* the node set (|*V*| = *n*), *E* the directed hyperedge set (|*E*| = *m*), *W* the weight set, and *f* a map from *E* onto *W*; a weighted directed hyperedge *e* (see **Figure 1a** for an example) is an ordered pair 〈*L, R*〉, with *L, R* ⊆ *V* and 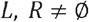, whose weight is *f*(*e*) = *w_e_* ∈ *W*; Nodes *v_i_* (*i* = 1,2,⋯, *n*) in *L* and *R* are called the tails and the heads of *e*, denoted as tails(*e*) and heads(*e*) and nodes(*e*) = tails(*e*) ∪ heads(*e*). Each tail or head has its own weight *w_j_* in the hyperedge, i.e., *v_j_* ∈ *L* ∪ *R*, and *w_e_* = ∑_*j*∈*R*_*w_j_* ×∑_*j*∈*R*_*w_j_*. A backward hyperedge, or simply B-edge, is a hyperedge 〈*L,R*〉 with |*R*| = 1 (**Figure 1c**). A forward hyperedge, or simply F-edge, is a hyperedge 〈*L,R*〉 with |*L*| = 1 (**Figure 1d**).

**Figure 1.**
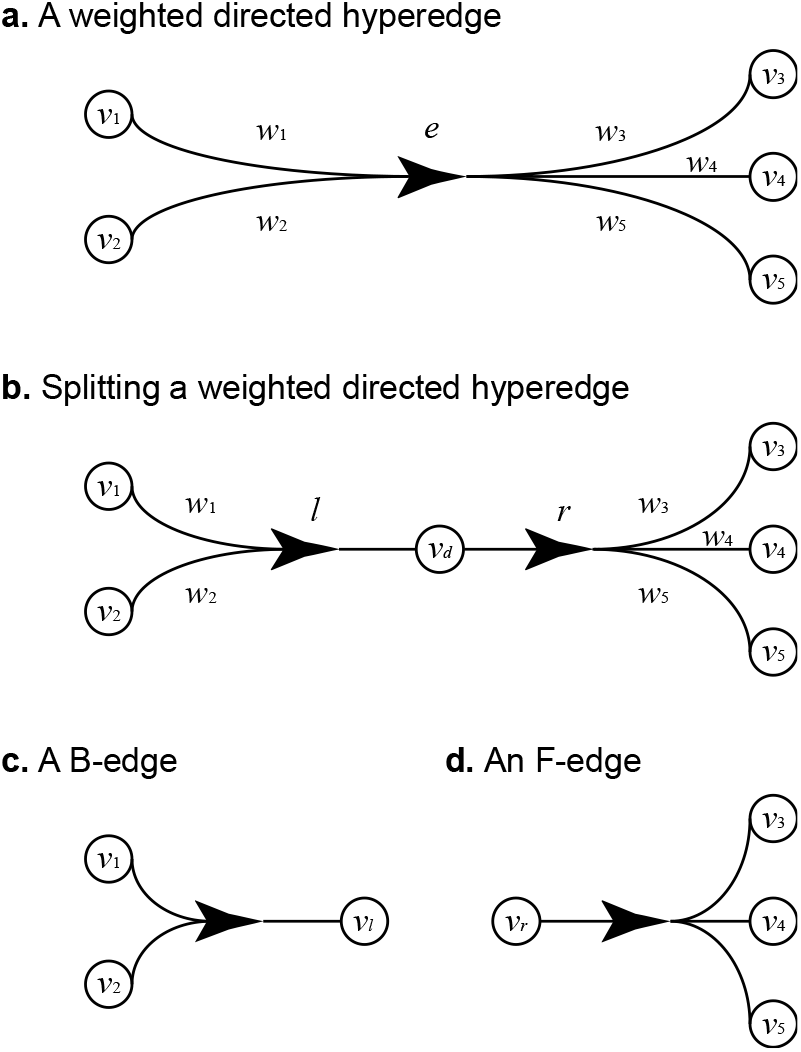
The relationship between a weighted directed hyperedge and a B-edge-F-edge pair, the directionality of the arrow represents the directionality of the hyperedge. **a.** A weighted directed hypergraph with one hyperedge *e.* **b.** *e* can be replaced by a B-edge *l* and an F-edge *r* which have a common node *v_d_*. **c.** A B-edge *l* generated by dividing node *v_d_* into two parts. **d.** An F-edge *r* generated by dividing node *v_d_* into two parts.

It is noteworthy that, as shown in **Figure 1**, a directed hyperedge e can always be replaced by a B-edge and an F-edge, through the addition of a dummy node *v_d_*, whose weight *w_d_* = 1, to the hyperedge (**Figure 1b**), *v_d_* is further decomposed into two dummy nodes *v_l_* and *v_r_* where *w_l_* = *w_r_* = l(**Figure 1c & d**). For the generated B-edge *l* and F-edge *r*, *f*(*l*) = ∑_*j*∈*L*_ *w_j_* × ∑_*j*∈*L*_ *w_j_* and *f*(*r*) = ∑_*j*∈*L*_ *w_j_* × 1, therefore, *l* and *r* respectively can represent the head node set and tail node set of the corresponding hyperedge e. The direction of a hyperedge is from the B-edge to the F-edge, i.e., from sending node set *L* (tails) to the target node set *R* (heads).

The biological interpretation of these definitions is as follows. In a multicellular interaction network, different cell types are modelled as different nodes, hyperedges represent communication among cell types, edge directions are from the cell types expressing the ligand to the cell types expressing the cognate receptor, and edge weights refer to the communication strengths. Then B-edges and F-edges (resulting from transformation from the original hyperedges) equate to the gene expression profiles of ligands and receptors across cell types. Hence, the pair of the only head of a B-edge *v_l_* and the single tail of an F-edge *v_r_* can serve as a possible ligand-receptor interaction.

Since each hyperedge can be divided into a pair of B-edges and F-edges, a weighted directed hypergraph can be specified by a matrix triple (*L,R,P*). The *n*× *x* matrix *L* can be viewed as the incidence matrix of all induced B-edges: *L*[*i,j*] (0 < *i* ≤ *n*, 0 < *j* ≤ *x*) is the weight of node *v_i_* in the B-edge *l_j_, i.e., v_i_* ∈ tails(*e_j_*), otherwise *L*[*i,j*] = 0. Similarly, the *n*×*y* matrix *R* is the incidence matrix of all induced F-edges: *R*[*i,j*] (0 < *i* ≤ *n*, 0 < *j* ≤ *y*) is the weight of node *v_i_* in the F-edge *r_j_, i.e.*, *v_i_* ∈ heads(*e_j_*) otherwise *R*[*i,j*] = 0. The *x*×*y* pair matrix *P*[*i,j*] (0 < *i* ≤ *x*, 0 < *j* ≤ *y*) records a pair of B-edge and F-edge that can form a weighted directed hyperedge in the directed hypergraph’ with: *R*[*i,j*] = 1 if B-edge *l_i_* and the F-edge *r_j_* can bind together, otherwise *P*[*i,j*] = 0. As an example, the matrix triple (*L,R,P*) in **Table 1** specifies the hypergraph in **Figure 1a**.

**Table 1.**
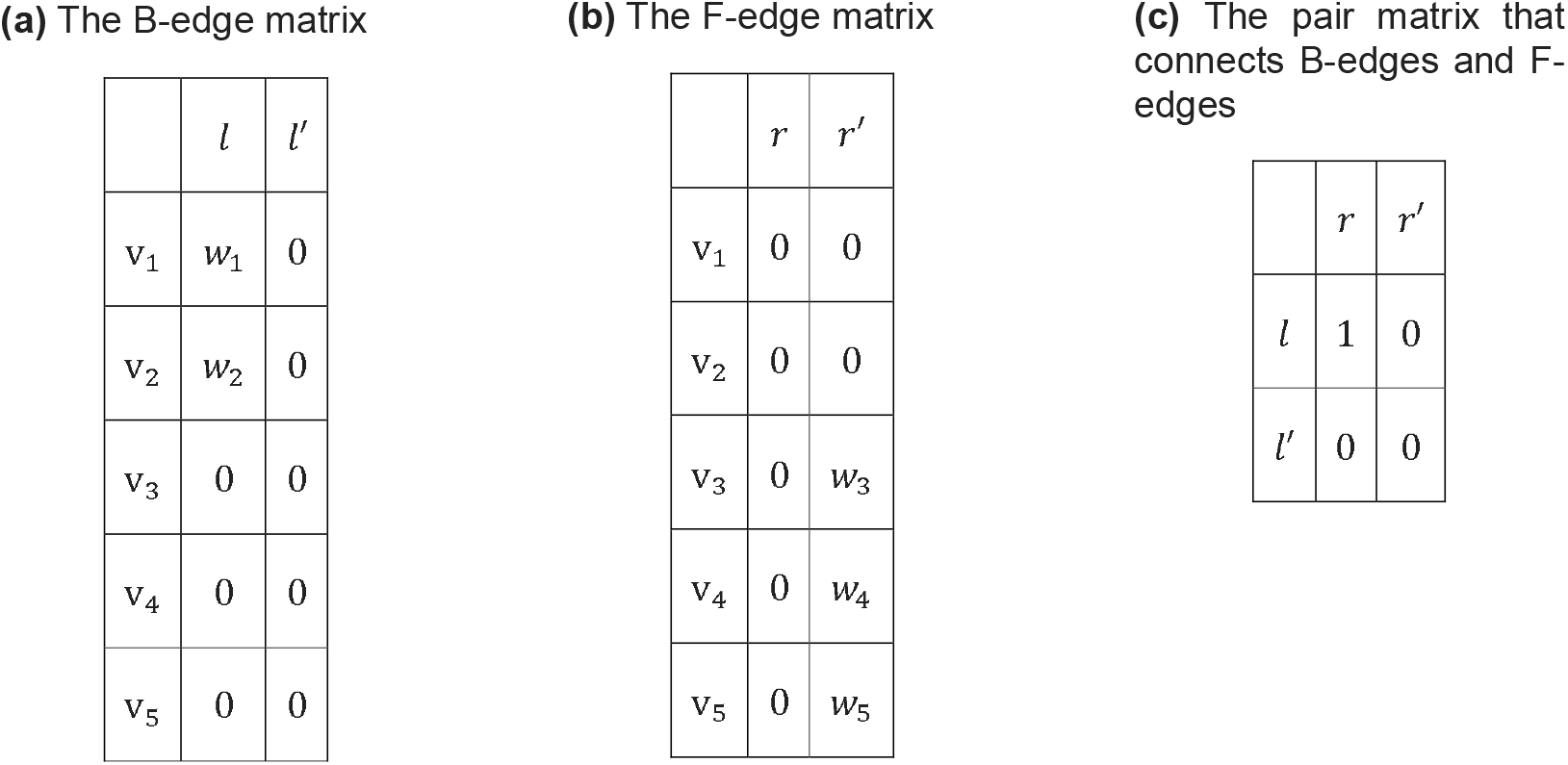
The triple matrices of a weighted-directed hypergraph.

**(a) The B-edge matrix**
| | $l$ | $l'$ |
| --- | --- | --- |
| $v_1$ | $w_1$ | 0 |
| $v_2$ | $w_2$ | 0 |
| $v_3$ | 0 | 0 |
| $v_4$ | 0 | 0 |
| $v_5$ | 0 | 0 |

**(b) The F-edge matrix**
| | $r$ | $r'$ |
| --- | --- | --- |
| $v_1$ | 0 | 0 |
| $v_2$ | 0 | 0 |
| $v_3$ | 0 | $w_3$ |
| $v_4$ | 0 | $w_4$ |
| $v_5$ | 0 | $w_5$ |

**(c) The pair matrix that connects B-edges and F-edges**
| | $r$ | $r'$ |
| --- | --- | --- |
| $l$ | 1 | 0 |
| $l'$ | 0 | 0 |

Because the unweighted directed hypergraph, which can be considered as a weighted directed hypergraph with uniform weights, is the best model to represent many-to-many relationships, it has been widely used in various areas [28–32]. Strongly-connected components in a directed hypergraph are components that can be replaced by single nodes [33]. To the best of our knowledge, there is not a definition of the densely connected subgraph, which cannot be collapsed into a single node, in a directed hypergraph. Here, we extend the definition of the density-based community structure from directed graphs to directed hypergraphs. A node community can be considered as a set of nodes that have more hyperedges within the node set than the other nodes. This is made precise in the following definition.

#### Definition 2.

(Community structure) A node community in a directed hypergraph is a non-empty node set *C* ⊆ *V* that satisfies one condition, which cannot be extended to any of its supersets (*C* ⊊ *S* ⊆ *V*) : |*E_C_*| > |*E*_O_|, where *E_c_* = {*e_c_*| nodes(*e_c_*) ⊆ *C*} and 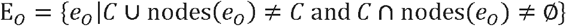. The edge set *E_C_* forms the incident hyperedge community of node community *C.*

In this paper, a node community is used to model a set of cell types that coordinate a biological process. From **Definition 2**, in a directed hypergraph, a set of hyperedges defines a group of densely connected nodes and forms a hyperedge community, which can be considered as a group of specific ligand-receptor pairs that connect multiple cell types in the cell type community and convey certain messages to regulate the biological process the cell type community involved. Hyperpaths [34, 35] and many other structures [36, 37] in the directed hypergraph impose constraints on the relationship between two adjacent hyperedges, *e.g.,* many heads of a hyperedge should be the tails of another hyperedge. On the contrary, there are no restrictions on any two directed hyperedges in a hyperedge community. This is because the regulatory relationship between two hyperedges is uncertain in many contexts. For example, within real world cell-to-cell communication networks, once a ligand binds to the target cell’s receptor and alters the target cell’s state, the target cell may change the activity of another ligand or receptor to initiate or inhibit another ligand-receptor-mediated communication [38, 39]. **Figure 2a** demonstrates a directed hypergraph with two node communities and two incident hyperedge communities.

**Figure 2.**
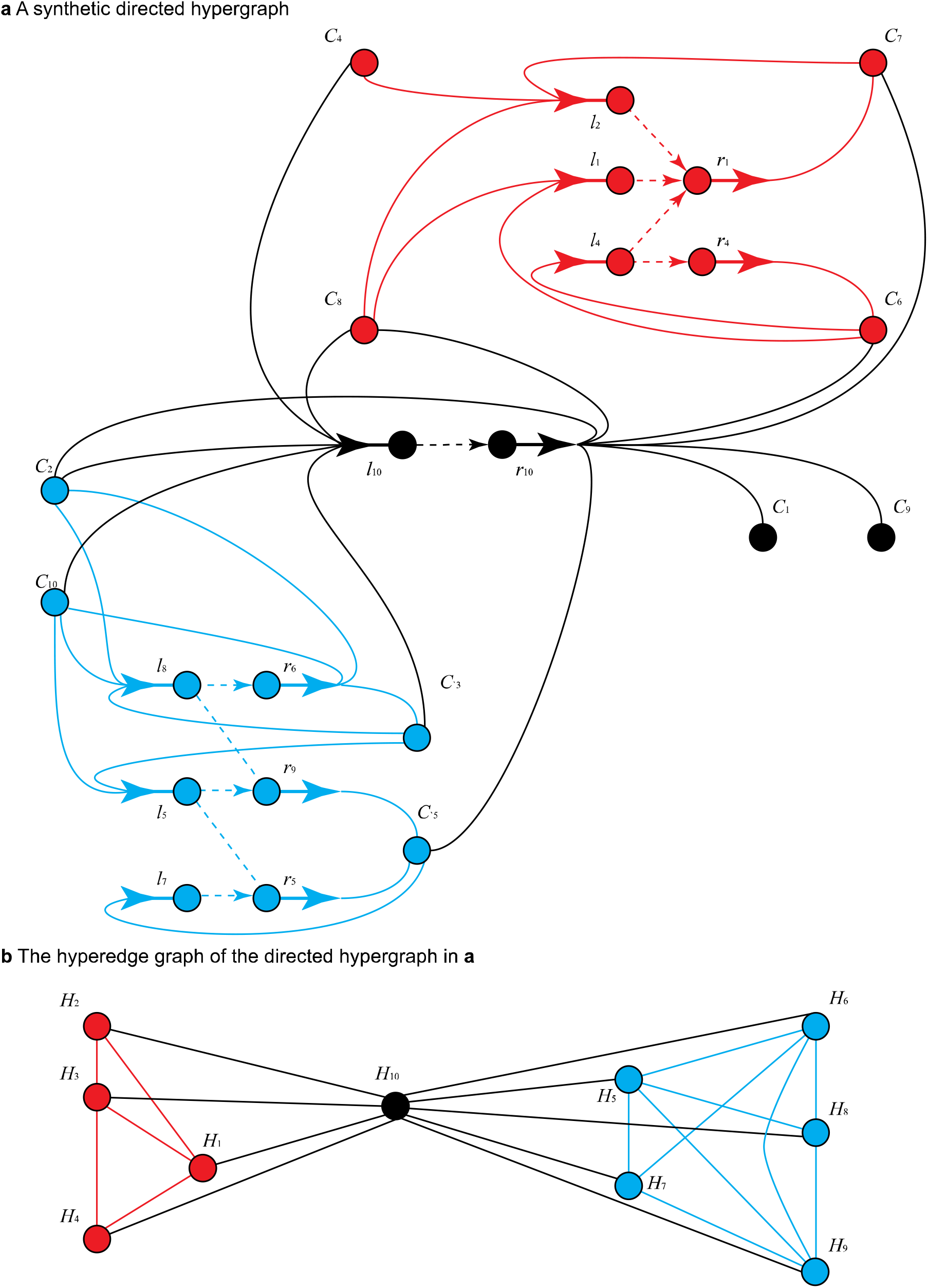
A synthetic directed hypergraph with ten nodes, ten hyperedges and two embedded communities. In this directed hypergraph, node IDs which start with *l* or *r* represent the derived dummy nodes (ligands and receptors), a directed hyperedge is represented by a B-edge and an F-edge. Solid lines connect nodes to construct a B-edge or an F-edge, dashed lined connect a B-edge and an F-edge to display a complete hyperedge (a ligand-receptor-mediated communication). Arrow directions indicate the hyperedge direction (from sending nodes to target nodes). Black nodes are outside of communities, and black hyperedges are housekeeping hyperedges. Other nodes and hyperedges of the same colour (red or blue) form a community structure. **a.** The visualisation of the synthetic directed hypergraph. **b.** The hyperedge graph of the synthetic directed hypergraph.

A feature of the hyperedges in a community is that they only connect a small subset of nodes within the entire network *i.e.* they have high specificities, highlighted in red and blue in **Figure 2a**. In the cell-to-cell communication network, the specificity of signalling via ligand-receptor pairs can be broadly viewed between two extremes. Housekeeping ligands and receptors are expressed in most, if not all, cell types at similar levels, thus all cell types in the network are connected by these signalling factors. Conversely, cell type-restricted ligands and receptors are only utilized by a relatively small portion of cell types making the network not evenly connected. In the corresponding hypergraph, specific hyperedges (red and blue hyperedges in **Figure 2a**) only connect a few nodes and form community structures while housekeeping hyperedges keep the whole hypergraph connected (black hyperedges in **Figure 2a**).

### 2.2 Community Detection Algorithms for Weighted Directed Hypergraphs

With the formal definition of a community structure defined above we next developed exact and approximate algorithms to detect these clustered nodes and hyperedges in the directed hypergraph. Ultimately, when applied to cell-to-cell communication networks, these methods aim to identify closely interconnected cell types (nodes) and associated intercellular signalling pathways (hyperedges) through the directed hypergraph model.

#### 2.2.1 Exact Detection Algorithms for Communities in the Weighted Directed Hypergraphs and Its Computational Complexity

To find all communities, all possible node sets in the directed hypergraph should be enumerated. For each of the node sets, both the number of hyperedges in the induced subgraph and the number of hyperedges connecting the subgraph and the rest of the hypergraph are used to examine if the induced subgraph is a community. In this way, the exact algorithm makes an exhaustive search to find all qualified communities in the hypergraph based on the definition. It is well known that community detection in the simple graph is NP-complete [40], which is equivalent to the problem of detecting communities in the directed hypergraph. Since it is infeasible using the exact algorithm to address this computationally intractable problem, we propose two approximate methods to find community structures in the directed hypergraph.

#### 2.2.2 Approximate Detection Algorithms for Communities in the Weighted Directed Hypergraphs

For simple graphs, the Markov Cluster algorithm (MCL) [41] and Louvain method [42] are two approximate clustering algorithms which have been widely used for community detection due to their competitive results and low time complexities. However, neither of them can be applied to directed hypergraphs without modification. To make them applicable for the hypergraph model, a directed hypergraph should be transformed to a simple graph first. Here, we propose a transformation approach from a directed hypergraph to a simple graph (specifically a hyperedge graph).

Although the relationships between nodes are many-to-many in the directed hypergraph, two interlocked hyperedges have a one-to-one mapping with each other. The weight of such a mapping is decided by the common nodes shared by two directed hyperedges. Since a node community is connected by a group of specific hyperedges and these hyperedges only connect the nodes within the community, hyperedges in a hyperedge community are more likely to interlock with each other than hyperedges outside of the community. Thus, if the relationships between directed hyperedges are modelled as simple edges and original directed hyperedges are modelled as nodes, hyperedge communities form many node communities. In this way, node community detection in a directed hypergraph is equivalent to hyperedge community identification in a hyperedge graph, in which hyperedges are nodes and two nodes are connected if the corresponding hyperedges have some common nodes in the original directed hypergraph.

#### Definition 3.

(Hyperedge graph) A hyperedge graph of a directed hypergraph, denoted as *G*(*C*), is a weighted simple graph such that each node *v_i_* in *V* (|*V*| = *m*) represents a hyperedge in the corresponding hypergraph *C*(*V, E, W, f*), two nodes *v_i_* and *v_j_* in *C* are connected by a weighted edge *e* if and only if their corresponding hyperedges share some common nodes in *C*. For a weighted directed hyperedge, the weight of *e* is the sum of the products of the weights of each common node in two hyperedges.

As mentioned before, a ligand-receptor binding may affect another ligand-receptor-mediated signalling pathway via a common cell (*e.g.* cell A may affect cell C, via cell B where cell A produces a ligand that binds a receptor on cell B, this binding regulates expression of a ligand from cell B that binds a receptor on cell C which regulates the state of cell C). The effect of one ligand-receptor pair on another pair is quantified by the product of the expression levels of the associated ligand and receptor that are expressed in the intermediate cell type. The overall effect through all shared cell types, *i.e.,* the weight of a simple edge in the hyperedge graph, is the sum of those products. **Figure 2b** illustrates the hyperedge graph transformed from the directed hypergraph in **Figure 2a**. Apparently, the hyperedges in those two communities form two independent hyperedge communities in the derived hyperedge graph. By obtaining closely connected hyperedge communities through a hyperedge graph, the incident node communities can be recognized. In this paper, we use the MCL and Louvain algorithms to identify hyperedge communities in the hyperedge graph, and the incident node communities in the related directed hypergraph.

## 3. Results

Previously, Ramilowski *et al.* [16] used messenger RNA (mRNA) as a proxy for protein expression and extracted expression levels of ligands and receptors in 144 primary cell types to construct the first cell-to-cell communication network in the human body. The underlying community structures in this network are unknown. In order to examine the versatility of the proposed community detection algorithms, we first apply them to synthetic directed hypergraphs embedded with predefined communities to assess their performance. Using the best detection strategy, we then applied it to two organism-wide datasets and identified many communities within.

### 3.1 Construction of directed hypergraphs with embedded communities

To ensure our synthetic hypergraphs shared characteristics of real cell-to-cell communication networks we first examined the expression patterns of ligands and receptors and the predicted connectivity in the high-quality human primary cell type atlas from Ramilowski *et al.* [16]. This revealed that the number of cell types expressing each ligand and receptor varied greatly. 60.6% of signalling factors (ligands and receptors) were broadly expressed across more than half the cell types, while 11.2% of signalling factors were specifically expressed in less than 10% of cell types (**Figure 3a**). From this analysis we identified four kinds of expression patterns for a signalling factor: i) the signalling factor is broadly expressed and its expression levels follow the uniform distribution (**Figure 3b** and **Supplementary Figure 1a**); ii) the signalling factor is broadly expressed and its expression levels vary greatly and tend to fit the exponential distribution (**Figure 3c** and **Supplementary Figure 1b**); iii) the signalling factor is only expressed in a small portion of the dataset and its expression levels have a uniform distribution (**Figure 3d** and **Supplementary Figure 1c**) and iv) the signalling factor is specifically expressed and the corresponding expressions exhibit a power-law distribution (**Figure 3e** and **Supplementary Figure 1d**).

**Figure 3.**
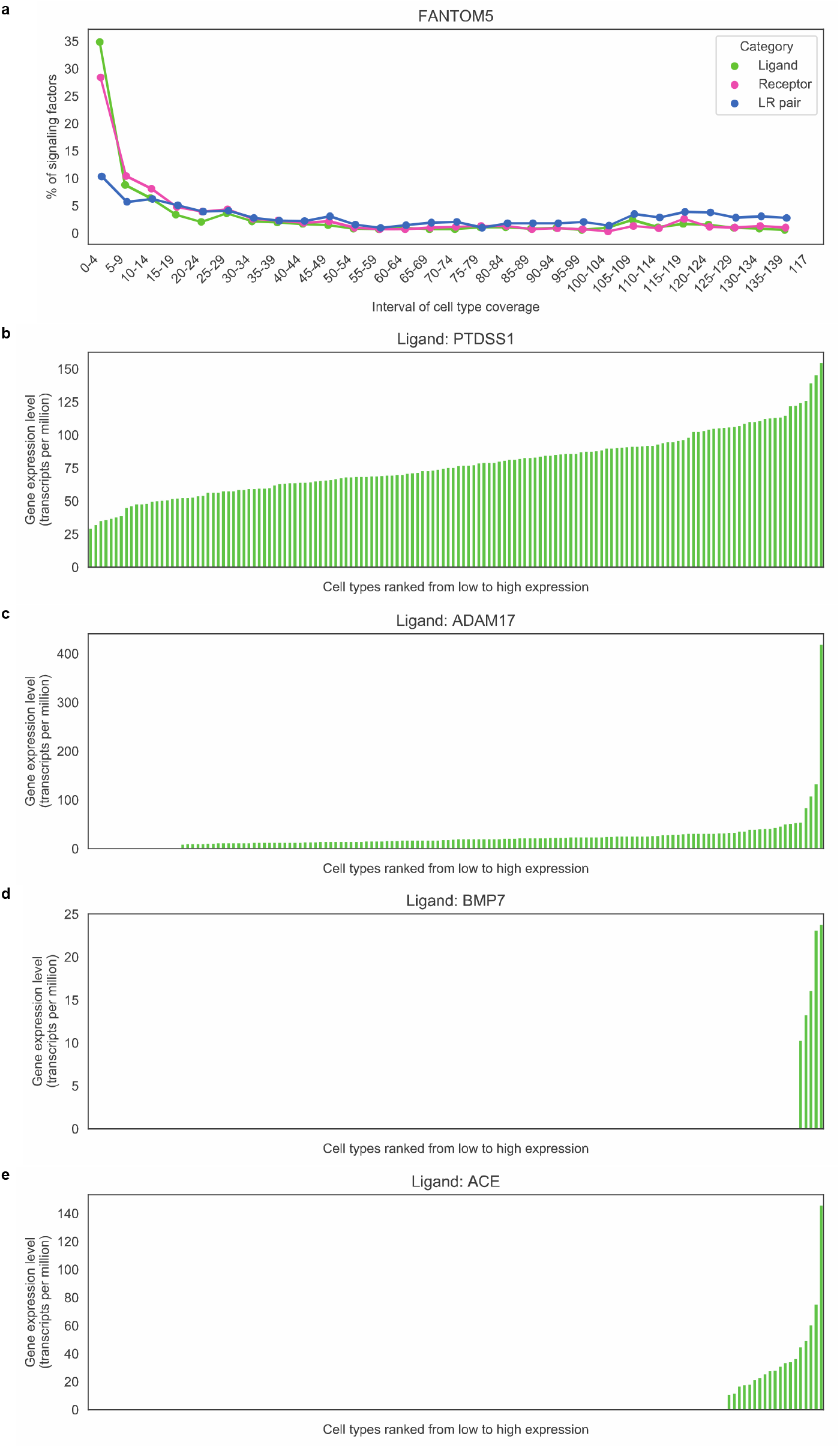
Gene expression patterns in the FANTOM5 dataset. **a.** The percentages of ligands (green), receptors (pink) only detected in certain amount of cell types and the proportions LR pairs that connecting a given number of cell types**. b-e.** The gene expression profile of a ligand in the FANTOM5 dataset. Ligands in **b** and **c** are broadly expressed while ligands in **d** and **e** are specifically expressed. Meanwhile, ligands in **b** and **d** have the uniform distribution when ligands in **c** and **e** follow the power law.

Based on the observations from this real data, we designed three kinds of weighted B-/F-edges i) housekeeping B-/F-edges, ii) weight-biased B-/F-edges and iii) specific B-/F-edges. These B-edges and F-edges were then combined to form several types of directed hyperedge. From **Figure 3b-e** and **Supplementary Figure 1**, housekeeping and weight-biased B-/F-edges are set to connect most nodes in the hypergraph while specific B-/F-edges are assumed to only connect nodes in the node community. Moreover, since the gene expression values of the ligands/receptors in the associated cell types can be processed in various methods to eliminate the sources of variability, node weights in a B-/F-edge do not directly stimulate the gene expression values from the obtained expression matrix, they are set to positive real numbers to simulate the normalized relative expression levels of the ligands/receptors without units and follow either power-law or uniform distribution. Note, node weights in the hyperedge communities of a directed hypergraph follow the identical distribution. Housekeeping hyperedges with similar values (less than 10) keep all nodes connected while the other hyperedges, whose minimum node weight is above 50, form hyperedge communities. If all hyperedge communities only have specific hyperedges, then the hypergraph is noted as a weight-specific directed hypergraph. Otherwise, the hypergraph is categorised as a weight-biased directed hypergraph because all hyperedge communities only have weight-biased directed hyperedges. In a weight-biased directed hyperedge, the weights of the nodes inside and outside of the node communities are taken randomly from two ranges. The weights of the nodes in the node community also follow a uniform or exponential distribution with a minimum value of 50, the weights of the other nodes are roughly the same and all less than 5.

After defining three kinds of directed hypergraphs, we constructed synthetic hypergraphs whose sizes (number of nodes) are 10, 20, 30, 40, 100, 200, 300 and 400. In order to test the performance of a community detection method, it is necessary to apply the method to hypergraphs with various settings. In terms of the size of a synthetic hypergraph, it should not be too small so that a community structure is hard to be embedded into the hypergraph. On the other hand, the hypergraph size should not be too large either to prevent excessive time consumption. Therefore, we varied graph sizes ranging between 10 to 400 nodes. In each hypergraph, the number of nodes and the number of directed hyperedges, which can be regarded as the pairs of distinct B-edges and F-edges, are equal. **Figure 2** shows a synthetic hypergraph with 10 nodes and 10 hyperedges. Furthermore, the number of embedded community structures in a hypergraph is proportional to the size of the directed hypergraph, such as 2, 4, 6, 8, 10, 20, 30 and 40. Since the directed hypergraph shown in **Figure 2** has 10 nodes and 10 hyperedges, two communities (red and blue communities) are embedded. Then 80% of the nodes and 96% of the directed hyperedges in the hypergraph are set to construct the community structures. Therefore, in **Figure 2**, 8 nodes are either red or blue, and 9 hyperedge are used to specifically connect these 8 nodes and form two hyperedge communities. Given the amounts of nodes, hyperedges and communities in a directed hypergraph, nodes and hyperedges are divided into certain number of groups to form communities. To evaluate the performance of a community detection algorithm in more general contexts, we fitted the community sizes (*i.e.,* number of nodes) and densities (*i.e*., ratio of hyperedges to nodes in a community) to follow uniform or exponential distribution. The construction of a synthetic directed hypergraph has four steps. First, the amounts of nodes and hyperedges inside and outside of the communities are calculated. Those nodes that belong to communities are randomly divided and assigned to communities so that the number of nodes in each community follows the given distribution. **Table 2** demonstrates how 40 nodes in a synthetic hypergraph are divided to follow certain community size distributions. The hyperedges inside of communities are then partitioned into the same amount of groups, the number of hyperedges in each partition is adjusted for several iterations until the ratio of the hyperedges to the nodes in every community is distributed uniformly or otherwise. **Table 3** shows the partitions of 40 hyperedges that are combined with node partitions in **Table 2** to satisfy the constraints of community density. Lastly, all hyperedges are randomly connected to the nodes based on the predefined rules, the weights of nodes in each hyperedge are also randomly allocated to match the given distribution. For a combination of directed hyperedge and community settings, we generated ten directed hypergraphs independently. These hypergraphs are then used to test the robustness of a community detection method.

**Table 2.** Examples of nodes partitions in a directed hypergraph with diverse community configurations. For 40 nodes, 35 of them are assigned to 8 node communities. Four kinds of hypergraphs are constructed: **a**) community size and density both follow the uniform distribution, **b**) community size follows the uniform distribution while community density follows the exponential distribution, **c**) community size follows the exponential distribution while community density follows the uniform distribution and **d**) community size and density both follow the exponential distribution.

|  | Community size:<br>uniform,<br>community<br>density: uniform | Community size:<br>uniform,<br>community<br>density: power | Community size:<br>power,<br>community<br>density: uniform | Community size:<br>power,<br>community<br>density: power |
| --- | --- | --- | --- | --- |
| Community 1 | 3 | 2 | 2 | 3 |
| Community 2 | 3 | 3 | 3 | 3 |
| Community 3 | 3 | 4 | 4 | 3 |
| Community 4 | 4 | 4 | 4 | 3 |
| Community 5 | 5 | 4 | 4 | 3 |
| Community 6 | 5 | 5 | 4 | 3 |
| Community 7 | 6 | 6 | 7 | 6 |
| Community 8 | 6 | 7 | 7 | 11 |
| Other nodes | 5 | 5 | 5 | 5 |

**Table 3.** Examples of hyperedge partitions in a directed hypergraph with diverse community configurations. For 40 hyperedges, 39 of them belong to 8 hyperedge communities. These 39 hyperedges are partitioned and combined with 8 node communities above to reach four community configurations: **a**) community size and density both follow the uniform distribution, **b**) community size follows the uniform distribution while community density follows the exponential distribution, **c**) community size follows the exponential distribution while community density follows the uniform distribution and **d**) community size and density both follow the exponential distribution.

|  | Community size:<br>uniform,<br>community<br>density: uniform | Community size:<br>uniform,<br>community<br>density: power | Community size:<br>power,<br>community<br>density: uniform | Community size:<br>power,<br>community<br>density: power |
| --- | --- | --- | --- | --- |
| Community 1 | 3 | 2 | 2 | 2 |
| Community 2 | 4 | 3 | 2 | 2 |
| Community 3 | 4 | 4 | 3 | 3 |
| Community 4 | 4 | 4 | 3 | 3 |
| Community 5 | 4 | 5 | 4 | 4 |
| Community 6 | 6 | 6 | 5 | 5 |
| Community 7 | 7 | 7 | 7 | 7 |
| Community 8 | 7 | 8 | 13 | 13 |
| Housekeeping<br>hyperedges | 1 | 1 | 1 | 1 |

In addition to the benchmark hypergraphs, fashioning appropriate metrics are also critical in determining the performance of a community detection algorithm. There is no universally accepted metric for this task, different metrics focus on various aspects of the problem, and are thereby biased to different approaches. In this paper, we denote the detection sensitivity as the fraction of the number of embedded communities entirely detected by our methods over the total number of embedded communities. Since the total number of node and hyperedge communities in a real cell-to-cell communication network is hard to estimate, we are interested in to what extent can we trust that the detected communities are true communities in the directed hypergraph. The detection precision, which is the ratio of the number of embedded communities entirely detected by our methods to the total number of detected communities that may partially overlap with the predefined communities, is used to examine this issue.

### 3.2 Community Detection in the synthetic directed hypergraphs

As described above, the synthetic directed hypergraphs were first transformed into hyperedge graphs, then MCL and Louvain community detection methods were applied to the hyperedge graphs to reveal the hyperedge communities. In each hyperedge community, the heads and tails of these hyperedges form the associated node community. As true cell-to-cell communication networks are weighted, we focus here on the weighted networks. For the weighted networks we modelled two scenarios: i) networks where edges were specific and weighted, and ii) networks where edges were broadly used but weighted with biases towards specific nodes.

By applying the two community detection methods to weighted hypergraphs, we found that MCL outperformed Louvain for hyperedge community detection for all graph sizes except the smallest of 10 nodes, and with larger graph sizes MCL’s performance improved while Louvain’s got worse (**Figure 4a&c**). However, for the weight-biased hypergraphs, the predicted node communities were larger than the actual node community (because the weight-biased hyperedges connect nodes inside and outside of the node communities and the community detection methods cannot distinguish these two kinds of nodes in a hyperedge). As a result, neither of these two methods can identify any predefined node communities in the weight-biased hypergraph. However, for the weight-specific hypergraphs, both of the methods have higher detection sensitivity and precision for node communities than those for hyperedge communities (**Figure 4a**). We also observed that the sensitivity of hyperedge community detection is lower than that of node community detection. This implies some correctly identified node communities were inferred from the wrongly detected hyperedge communities. These wrongly detected hyperedge communities must be subsets of the true hyperedge communities because specific hyperedges are designed to only connect the nodes within a node community while housekeeping hyperedges connect other nodes which makes the inferred node communities bigger than the predefined node communities.

**Figure 4.**
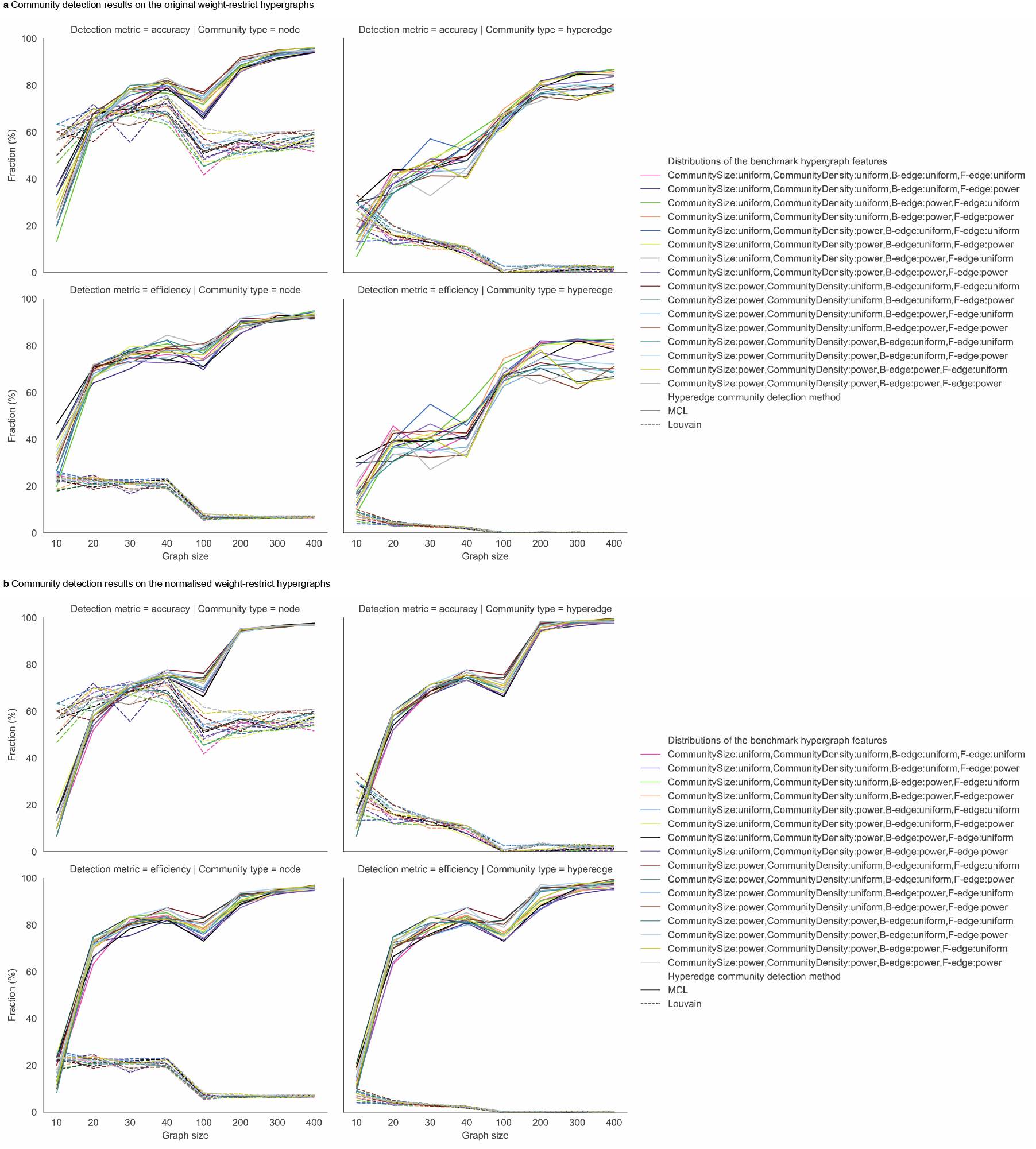
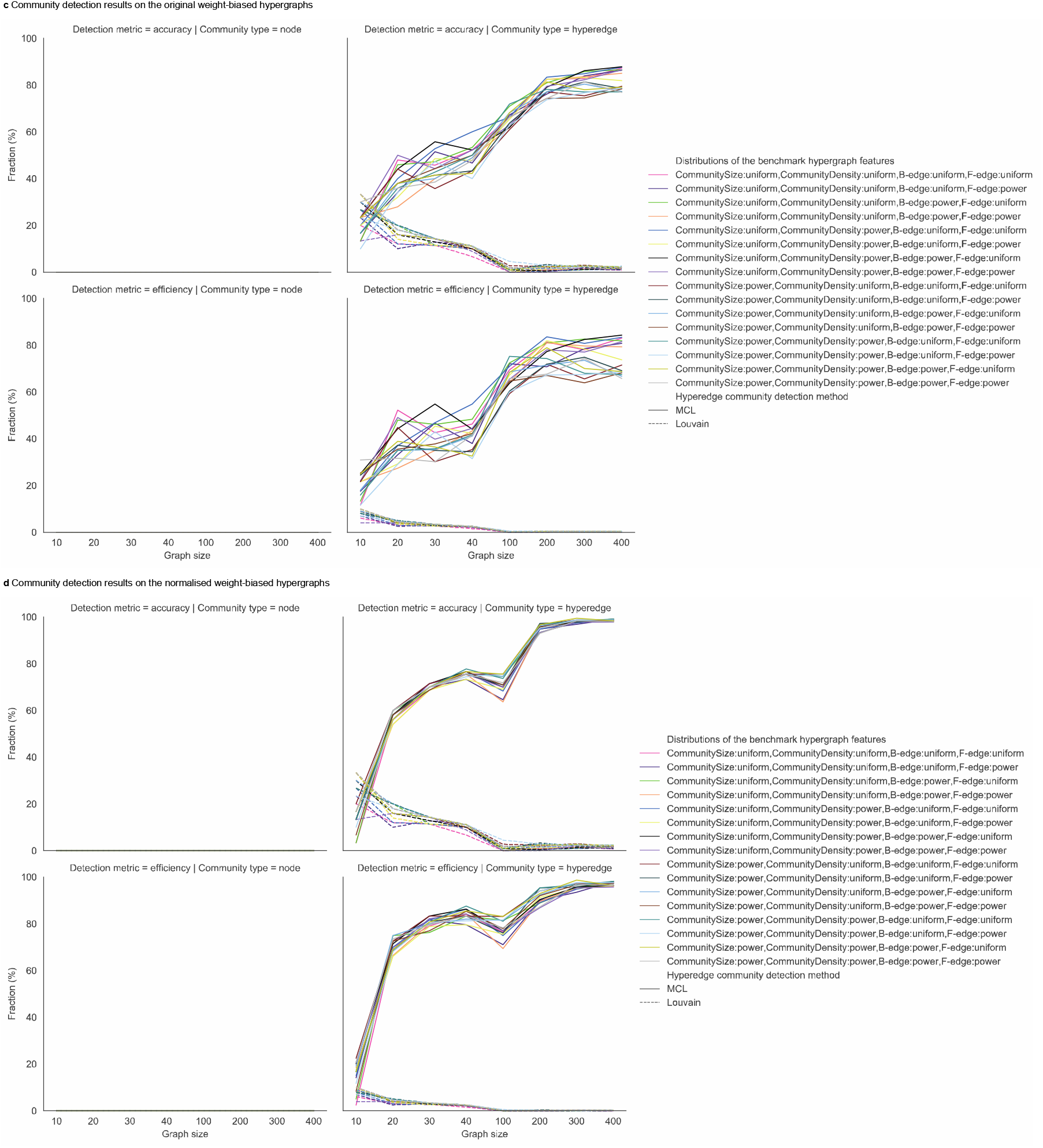
Community detection results on weighted directed hypergraphs using MCL-based method (solid lines) and Louvain-based method (dashed lines). Both MCL and Louvain are applied to graphs across a range of sizes (10 to 400 nodes) and with variable community sizes and community density. Community size (number of the nodes that are designed to be inside of a community), community density (the ratio of directed hyperedge amount within each hyperedge communities to the number nodes in the corresponding node community), node weights in a B-edge or a F-edge are uniformly or power-law distributed. Left columns show detection sensitivity (the fraction of detected true communities over all embedded communities) and precision of node communities (the ratio of the detected communities to all detected communities), while the same metrics for hyperedge communities are presented on the right. **a** and **b** show the results on weighted-specific hypergraphs whereas **c** and **d** display the results on weighted-biased hypergraphs. Node weights in **a** and **b** are normalised when they were used in **c** and **d**.

In a hyperedge graph, hyperedges with higher weights have a higher influence on the results than those with lower weights. To minimize such discrepancies, node weights can be normalised so the sum of heads/tails in a directed hyperedge is one (noted as specificity) and all hyperedges have identical weights. For the weighted directed hypergraphs, hyperedge and node community detection was improved through normalisation, however, this did not improve node community detection in the weight-biased hypergraphs (**Figure 4b&d**). Overall, the MCL-based method is more likely to identify predefined node and hyperedge communities in the weighted directed hypergraphs with diverse settings.

### 3.3 Community Detection in weighted directed hypergraphs based on real datasets

In the previous section, we have shown that using the MCL-based method and node weight normalisation can efficiently identify more underlying communities than the other community detection strategies. This strategy, collectively referred to as HyperCommunity algorithm in the following, is thus applied to directed hypergraphs generated from two organism-wide single-cell datasets using microwell-seq technology: the Human Cell Landscape (HCL) [43] and the Mouse Cell Atlas (MCA) [44]. There are 82 cell types in adult tissues from HCL and 88 cell types in adult tissues from MCA. To eliminate the noise within gene expression data, we required that the ligands and receptors considered needed to be detected in more than 10% of cells of at least one cell type, which is the detection threshold used in the original paper to perform receptor–ligand pairing analysis [45]. To construct the cell-to-cell communication network 2,293 human ligand-receptor pairs (connectomeDB2020) with primary literature support are used [46]. For analysis of the MCA dataset, 1,895 mouse ligand-receptor pair homologs of the human pairs were extracted through the NCBI HomoloGene Database [47] (**Supplementary Table 3**). For each dataset, analyses were carried out separately for cell-to-cell communication via cell-surface ligands (which require physical contact between the sending and target cells) and via secreted ligands (which can occur between two distant cells).

For ligand-receptor pairs using cell-surface ligands, 7.26% (9/124) ligands and 4.03% (6/149) receptors were detected in at least half of cell types in HCL, 7.29% (7/96) ligands and 3.15% (4/127) receptors were detected in at least half of cell types in MCA. For ligand-receptor pairs using secreted ligands, 6.34% (22/347) of ligands and 3.64% (11/302) of receptors were detected in at least half of cell types in HCL, 5.41% (17/314) of ligands and 2.84% (8/282) of receptors were detected in at least half of cell types in MCA. Hence, the weighted directed hypergraphs from these two real datasets have both weight-specific and weight-biased hyperedges. Based on our analysis in the previous section, these weight-biased hyperedges can affect the performance of node community detection.

Communities identified in the cell-to-cell communication networks based on plasma membrane and secreted ligands, using the MCL-based detection strategy with normalisation pre-processing, are summarised in **Supplementary Table 4**. 11 of the 19 predicted communities connected at least 90% of cell types (5 connected all cell types), which implies the underlying hyperedge (ligand-receptor pair) communities include broadly expressed ligands and/or receptors. We, therefore, next pruned the network to remove connections to cell types with specificity below 0.2, which effectively removes housekeeping hyperedges. As illustrated in **Supplementary Table 5**, this identified smaller communities that connected fewer cell types compared to in the unpruned networks (**Supplementary Table 4**). In the HCL dataset, 7 communities were identified based on plasma-membrane ligands (connecting 20, 4, 5, 15, 6, 33 and 25 cell types) and 8 based on secreted ligands (connecting 8, 51, 22, 3, 2, 59, 9, and 35 cell types), while in the MCA dataset, 7 communities were identified based on plasma-membrane ligands (connecting 36, 1, 17, 15, 19, 9 and 5 cell types) and 8 based on secreted ligands (connecting 13, 10, 17, 25, 14, 75, 26 and 61 cell types). Together this suggests pruning can help reduce the number of trivial predicted communities that connect all nodes. **Figure 5** shows a community, which was identified by HyperCommunity, based on secreted ligands in the MCA dataset. In this community, stromal cells can only send ligands to blood cells. While these blood cells are able to send and capture ligands from both stromal and blood cell.

**Figure 5.**
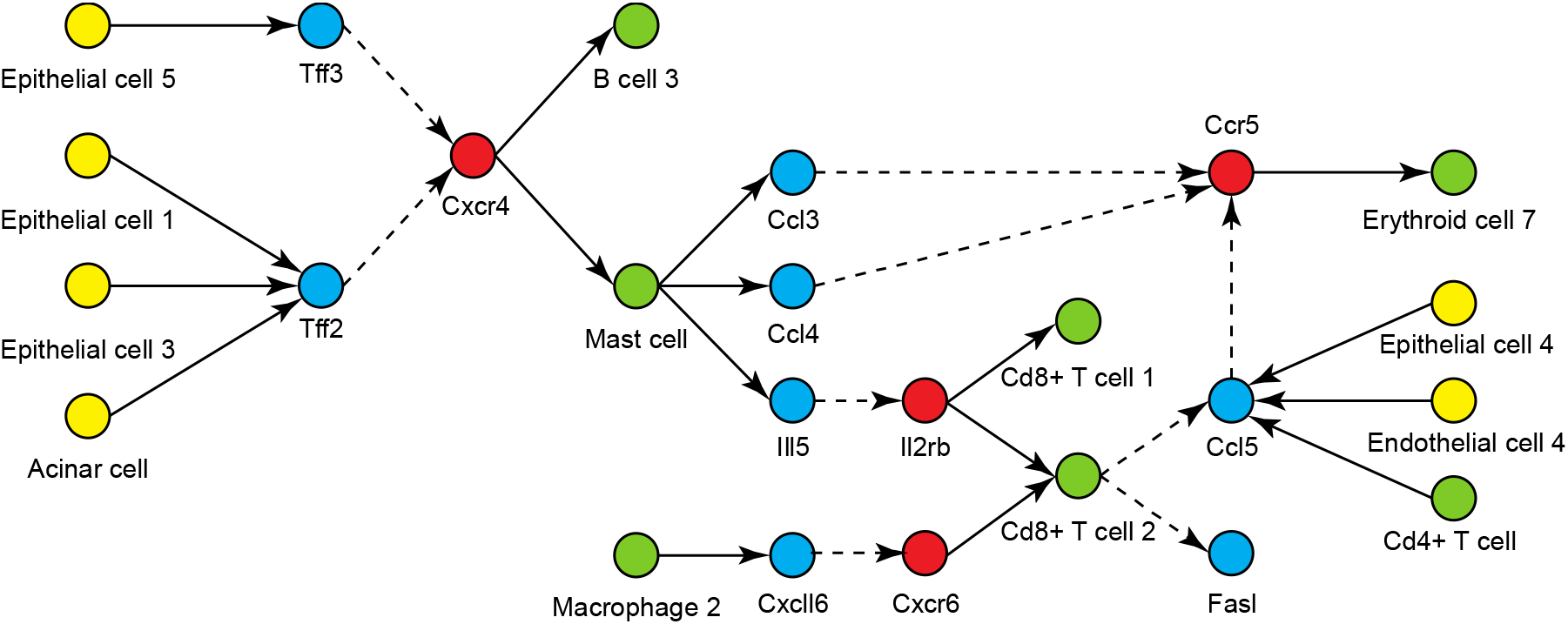
A secreted-ligand-mediated community identified in the MCA dataset. Yellow nodes are stromal/endothelial cells and green nodes are blood/immune cells. Those cell types expressing ligands are connected to the corresponding blue nodes while the cell types with receptors are connected to the related red nodes. A directed dashed edge connects a secreted ligand (blue node) to its cognate receptor (red node).

Of note, community detection using HyperCommunity identifies disjoint ligand-receptor pair communities, however, cell type communities tend to overlap with each other (see **Supplementary Figure 2**). Using PM ligands, a cell type can be involved in at most 5 communities in the HCL dataset and 4 communities in the MCA dataset, whereas, through secreted ligands, a cell type can participate at most in 6 communities in the HCL dataset and in 5 communities in the MCA dataset. Overall, HyperCommunity algorithm identified more cell type communities connected by secreted-ligand-mediated communication and these communities are more likely to be overlapped.

## 4. Discussion

Metazoans rely heavily on ligand-receptor-mediated intercellular communications to modulate biological events [5]. The maintenance of homeostasis depends on the proper orchestration of many cell types and various signalling pathways [3]. To capture the complex multicellular interactions and how the signals are transmitted through ligand-receptor axes, we used a new mathematical model, the weighted directed hypergraph, to model the many-to-many relationships among cells in a biological environment. We proposed the first definition of the community structure in a directed hypergraph and designed several algorithms to detect these node and hyperedge communities.

By applying two approximate detection strategies to the benchmark hypergraphs, we showed that the MCL-based algorithm could sensitively and precisely distinguish embedded communities in the weight-specific directed hypergraphs with different configurations and hyperedge communities in various weight-biased directed hypergraphs. We also showed that normalisation of node weights in the directed hyperedge yielded higher detection sensitivities and precisions. We then applied this best community identification method to gene expression data from organism-wide human and mouse datasets. Many disjoint ligand-receptor pair communities and overlapped cell type communities were identified. With a cell type community, specific ligand-receptor pairs connect cell types from distinct lineages together. Nevertheless, there are two limitations in the used real datasets. First, broadly expressed ligands and receptors hinder the identification of celltype and ligand-receptor communities thus they should be removed prior to analysis (otherwise need methods that down-weight these). The other limitation is that we have used the original cellular annotations provided by the original manuscripts [43, 44], if the annotations are incorrect or imprecise this will affect the observed results.

Some simple cell type communities have been observed. For example, germline stem cells and surrounding postmitotic somatic cells form a stem-cell niche in the Drosophila testis and establish a self-renewing signalling-mediated community [48]. Besides, individual tumour cells, which leave the primary tumour, spontaneously cluster together and induce collaborative interactions. These tumour cell communities have been linked to metastasis promotion [49]. Nevertheless, these investigated cell type communities are very simple and physically isolated. More complex communities are difficult to validate using current technologies. While this study cannot determine conclusively all exact cell type sets and ligand-receptor interaction groups, it does suggest that the cells in the body are not evenly connected. Through diverse ligand-receptor-mediated pathways, cell types arising from different lineages collectively orchestrate tissue homeostasis. With the availability of large amount of transcriptome data, more context-dependent cell type and ligand-receptor pair communities in the cell-to-cell communication network will be recognized through this weighted directed hypergraph model. In conclusion, we expect that revealing community structures in the weighted directed hypergraph model of a cell-to-cell communication network can illuminate the properties of biological systems that were previously unrecognized in the simple graph model, and facilitate the understanding of the underlying complexity of biology.

## Acknowledgements

We would like to acknowledge and thank Dr. Elena Denisenko for her insightful comments. This work was carried out with the support of a collaborative cancer research grant provided by the Cancer Research Trust ‘Enabling advanced single-cell cancer genomics in Western Australia’, a grant from Cancer Council of Western Australia and an Australian National Health and Medical Research Council project grant APP1146323. R.H. is supported by an Australian Government Research Training Program (RTP) Scholarship. A.R.R.F. was supported by funds raised by the MACA Ride to Conquer Cancer and a Senior Cancer Research Fellowship from the Cancer Research Trust. A.R.R.F. is currently supported by an Australian National Health and Medical Research Council Fellowship APP1154524. Analysis was made possible with computational resources provided by the Pawsey Supercomputing Centre with funding from the Australian Government and the Government of Western Australia.

## Supplementary Tables

**Supplementary Table1. Community detection results on original benchmark hypergraphs using the MCL-based and Louvain methods.**

**Supplementary Table 2. Community detection results on normalized benchmark hypergraphs using the MCL-based and Louvain methods.**

**Supplementary Table 3. The ligand-receptor pairs used to identify cell type communities and ligand-receptor pair communities in the HCL and MCA datasets.**

**Supplementary Table 4. Community detection results on the HCL and MCA datasets using all LR pairs.**

**Supplementary Table 5. Community detection results on the HCL and MCA datasets using all LR pairs when the normalised node weight threshold is 0.2.**

## Supplementary Figures

**Supplementary Figure 1.**
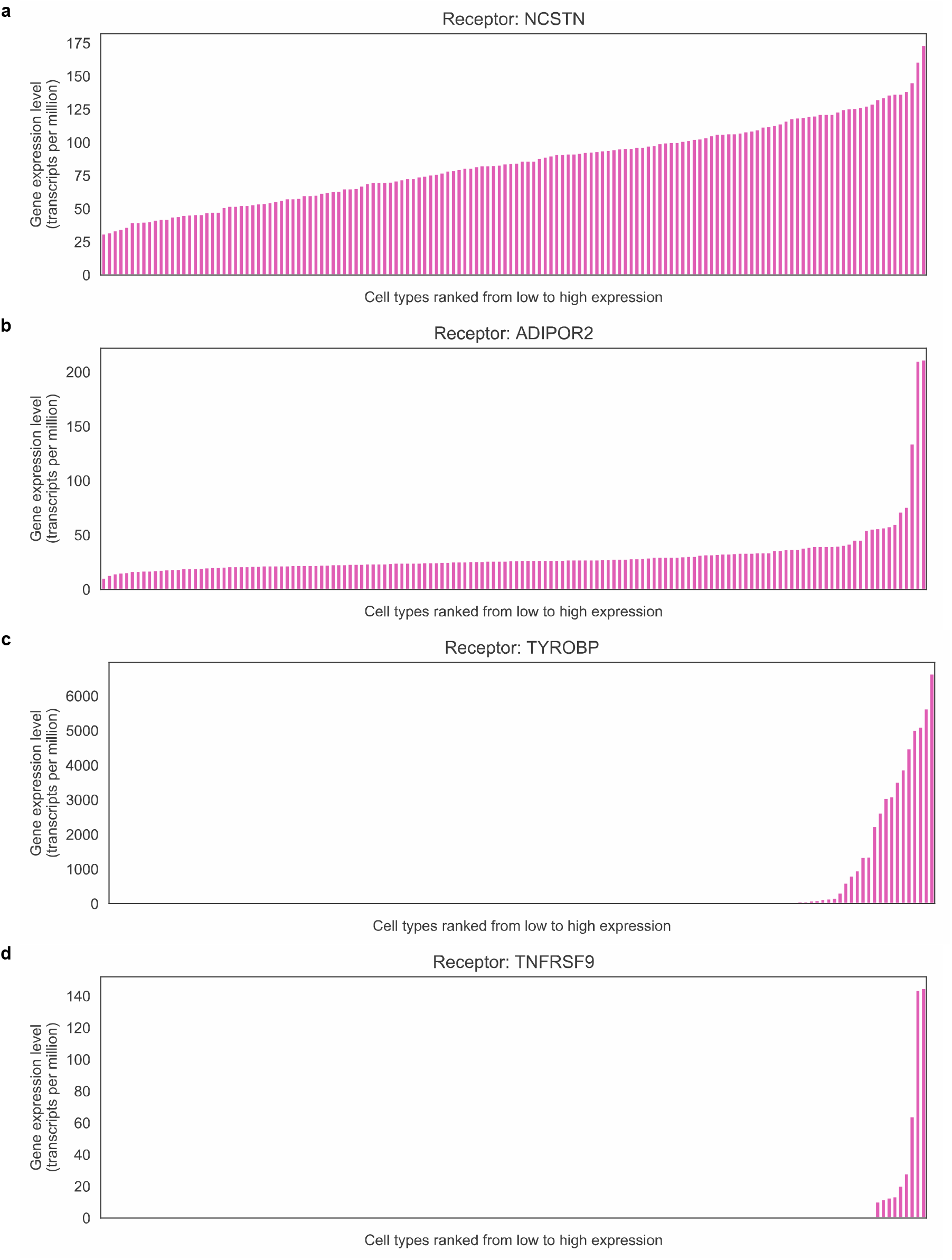
Receptor expression patterns in the FANTOM5 dataset. The gene expression profile of a receptor in the FANTOM5 dataset. Receptors in **a** and **b** are broadly expressed while receptors in **c** and **d** are specifically expressed. Meanwhile, receptors in **a** and **c** have the uniform distribution when receptors in **b** and **d** follow the power law.

**Supplementary Figure 2.**
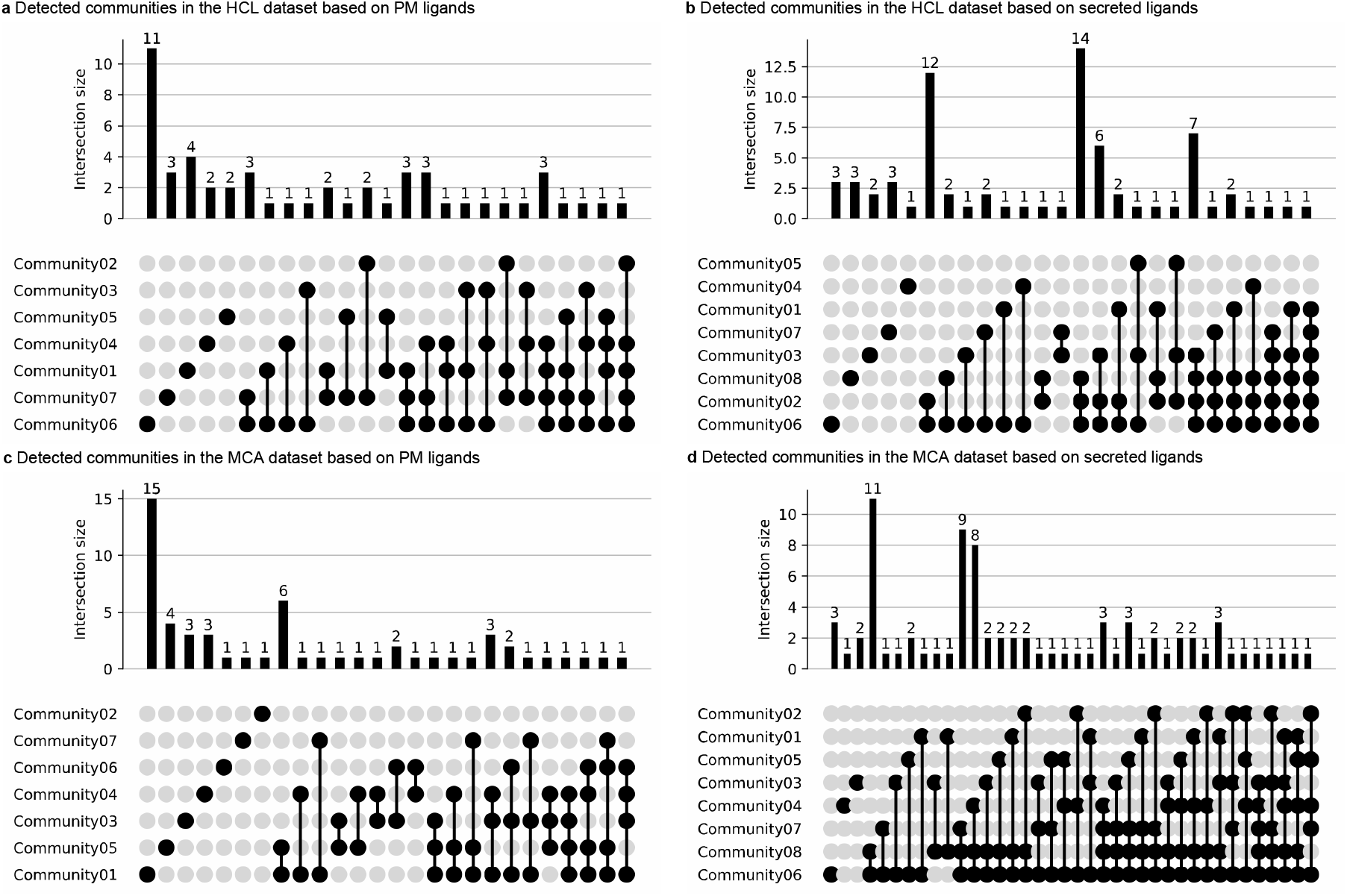
UpSet plots showing the intersection of detected communities in the HCL and MCA datasets using PM or secreted ligands. In the HCL dataset, HyperCommunity algorithm detected 7 communities based on PM ligands (**a**) and 8 communities based on secreted ligands (**b**). In the MCA dataset, HyperCommunity algorithm also detected 7 communities based on PM ligands (**c**) and 8 communities based on secreted ligands (**d**).

